# The determinants of aggression in male Siamese fighting fish, *Betta splendens*

**DOI:** 10.1101/2020.04.16.045179

**Authors:** R. E. McGoran, S. K. Papworth, S. J. Portugal

**Affiliations:** School of Biological Sciences, Royal Holloway University of London, Egham, Surrey, UK TW20 0EX

**Keywords:** Behaviour, Boldness, Male-male displays, Neophobia, Spectrophotometry, Territoriality

## Abstract

Siamese fighting fish, *Betta splendens*, have been extensively studied due to their aggression and stereotypical displays. Many studies have focused on their characteristic opercular flaring, while the less aggressive and less energetically costly lateral display have been comparatively understudied. Many factors have been shown to influence aggression in *Bettas*, notably body length and the personality trait of boldness, however, the role that colour plays in determining an individual’s aggressiveness is much less clear. The role of colour has only been briefly studied, and based on human interpretations of colour, i.e. limited to what the receivers’ eyes and sensory systems actually can process and discriminate, with results suggesting blue males are more aggressive than red males. Using male-male interactions, measuring opercular flaring and lateral display we found that colour and personality do play a role in determining the degree of aggressiveness in *Betta splendens*. Blue-finned males were more aggressive, performing longer lateral displays more frequently. Blue fins are a phenotype observed in wild type males and is likely selected for to allow visual cues to travel through the murky water they inhabit. Body mass was positively correlated with lateral display frequency, and opercular flare frequency and duration. Finally, neophobic individuals – individuals that were less willing to approach a novel object – were more aggressive, performing significantly more lateral displays. This indicates that personality may impact fighting strategy, with males either choosing to end conflicts quickly with more aggressive displays or to outlast their opponent with less energetically costly displays.

## INTRODUCTION

Animals compete aggressively with individuals of the same species over limited resources, such as territory and mates (Parker 1974). Individuals perform displays to settle disputes for resources, conveying information about their physiological condition, and their willingness to fight (Parker 1974). Successful displays can increase access to mates, mating success and territory size (Alton et al. 2013). Siamese fighting fish (*Betta splendens*) are an extreme example of an aggressive species. *B. splendens* are a facultatively air-breathing freshwater Anabantoid fish, which inhabit hypoxic waters where air-breathing is essential (Graham 1997). Domestic breeds have long been considered model organisms for behavioural studies due to their aggression, and stereotypic and conspicuous displays (Tate et al. 2017). Male *B. splendens* have multiple visual displays, including fin flaring and opercular flaring, which are designed to intimidate opponents by increasing their perceived body size (Simpson 1968). Examining social interactions in *B. splendens* has suggested that opercular flaring (raising of the operculae) and lateral displays (fin flaring while showing their broadside) are the most common aggressive displays in male-male and male-female interactions (Forsatkar et al. 2016). Displays often start with low-level aggressive behaviours, i.e. fin flaring, which can develop into more aggressive actions, i.e. opercular flaring and biting (Forsatkar et al. 2016). The duration of opercular flaring relates to an individual’s condition and has been demonstrated to be a reliable predictor of the winner of a male-male aggressive interaction (Simpson 1968).

Historically, much focus has been on opercular flare duration as an indicator of aggression (Tate et al. 2017). This study aims to explore what characteristics relate to an individual’s aggression by studying morphological and behavioural traits. Several characteristics have been linked to aggression in *B. splendens* including boldness (Hebert et al. 2014), colour (Simpson 1968) and size (Jaroensutasinee and Jaroensutasinee 2001). Subsequently, this study predicts that aggressive male *B. splendens* will display bolder personality traits (greater exploration and less neophobic in a novel environment), coupled with a larger body size. The threat displays of *B. splendens* are highly visual, thus it is likely colour contributes to aggression (Simpson 1968). Studies investigating the role of colour in animal behaviour have typically relied on categorical human assessments of colour, which are likely to underestimate or miscategorise the true colour of an individual. Therefore, using spectrophotometry, we determined the true colour of individual *B. splendens* and predict that those reflecting a shorter wavelength will be more aggressive.

## METHODS

### Subjects and housing

Adult male *B. splendens* (N =19) were housed individually in visually isolated 10.5 x 28 x 16.5 cm tanks and kept in laboratory conditions (light:dark cycle 14:10 h, 25 ± 1°C, 7.8–8.2 pH). Males were kept in isolation in their home tanks for at least 7 days prior to the behavioural observations to reduce the effects of prior social experience on their responses. Individuals were fed once a day, six days a week, on a mix of *Daphnia* spp., *Artemia* spp. and Tetra™ *Betta* flake food. The males were wet weighed to record their body mass (g) and their body and fin colour were recorded using spectrophotometry (see supplementary material for full methods).

### General experimental set-up

For all experiments, the tanks contained tap water (pH 7.4) aged for at least 18 h overnight at a temperature of 24.9 ± 0.96°C. Due to a limited supply of water from the home tank system aged tap water was used. All tanks were covered to remove external stimuli, except in the aggression trial where the front was left uncovered for observations. All behaviour was recorded using the PC event-recording software, Stopwatch+.

### Aggression

Randomly selected males were placed in two separate tanks (35 × 18 × 23.5 cm) placed together lengthwise with opaque paper preventing opponent observations. Males were given an hour to acclimatise before the partition was removed and their behaviour was recorded for 20 minutes. The following behaviours were recorded: (a) the duration (Od) and frequency (O_f_) of opercular flaring, (b) the duration (L_d_) and frequency (L_f_) of lateral displays. Each individual was observed once per day and repeat encounters were allowed to occur. Males met each opponent between 1 and 5 times. Male aggression was assessed using David’s score (DS, Gammell et al. 2003).

### Boldness

Boldness trials were performed in a 48 x 33 x 11 cm tank divided into thirds: a sheltered (containing a cave, plastic weeds and a plastic cover to create shade), intermediate (containing plastic weeds) and exposed zone (well-lit empty and shallow section with a predator silhouette hanging above). A randomly selected male was placed into the sheltered end and held in place by a Perspex screen. The fish were given five minutes to acclimatise before the screen was removed and the time spent in each section was recorded for 20 minutes. Individuals were tested once a week for three weeks. A boldness score was generated by weighting each section of the tank (see Portugal et al. 2017a). The sheltered, intermediate and exposed zones were weighted as one, two and three times the total length of time an individual spent in each respectively. The scores were averaged over three repeats.

### Neophobia

Neophobia trials were performed in the aggression tanks, which fish were already highly accustomed too, so that only introduced novel objects would influence behaviour. The tank was split into thirds. A novel object was placed in one end. The experimental design was identical to the boldness trials with regards to repeats, recording and weighting of the three zones. A different novel object was used for each trial.

### Statistical analysis

Repeatability values were calculated using a one-way ANOVA for boldness and neophobia, providing a measure of individual consistency. Spearman’s correlation was calculated between the repeatable boldness and neophobia traits for signs of any behaviours measuring the same trait. Any significantly correlated traits were reduced to the most repeatable measure. Linear models were performed to determine which variable related to aggression. All analysis was performed using R, version 3.3.1 (R Development Core Team, 2013).

## RESULTS

### Boldness and neophobia

Individual boldness was not consistent across trials (bold score: *R* = −0.05, *P* = 0.64; latency to leave the sheltered zone: *R* = 0.007, *P* = 0.46; latency to enter the exposed zone: *R* < 0.001, *P* = 0.48). However, individual neophobia was repeatable across trials (neophobia score: *R* = 0.45, *P* < 0.001; latency to enter the novel object zone: *R* = 0.47, *P* < 0.001). Both neophobia measures were significantly correlated (Spearman’s rank correlation: *r* = −0.835, *N* = 19, *P* < 0.001), hence, the latency to enter the novel object zone was used in the linear model.

### Colour

Average fin colour of the Bettas ranged from 482.7 to 608.5 nm, while the average body colour ranged from 472.9 to 604.8 nm. The visible light spectrum includes: violet (400-450 nm), blue (450-500 nm), green (500-570 nm), yellow (570-590 nm), orange (590-610 nm), and red (610-700 nm) light.

### Linear models

The linear models (DS L_d_, L_f_, O_d_, O_f_) accurately fitted the data (table 1). The models showed that individuals with bluer colouration and larger body size were more aggressive. Model DS Lf revealed that neophobic males were more aggressive (fig 1).

**Fig. 1.**
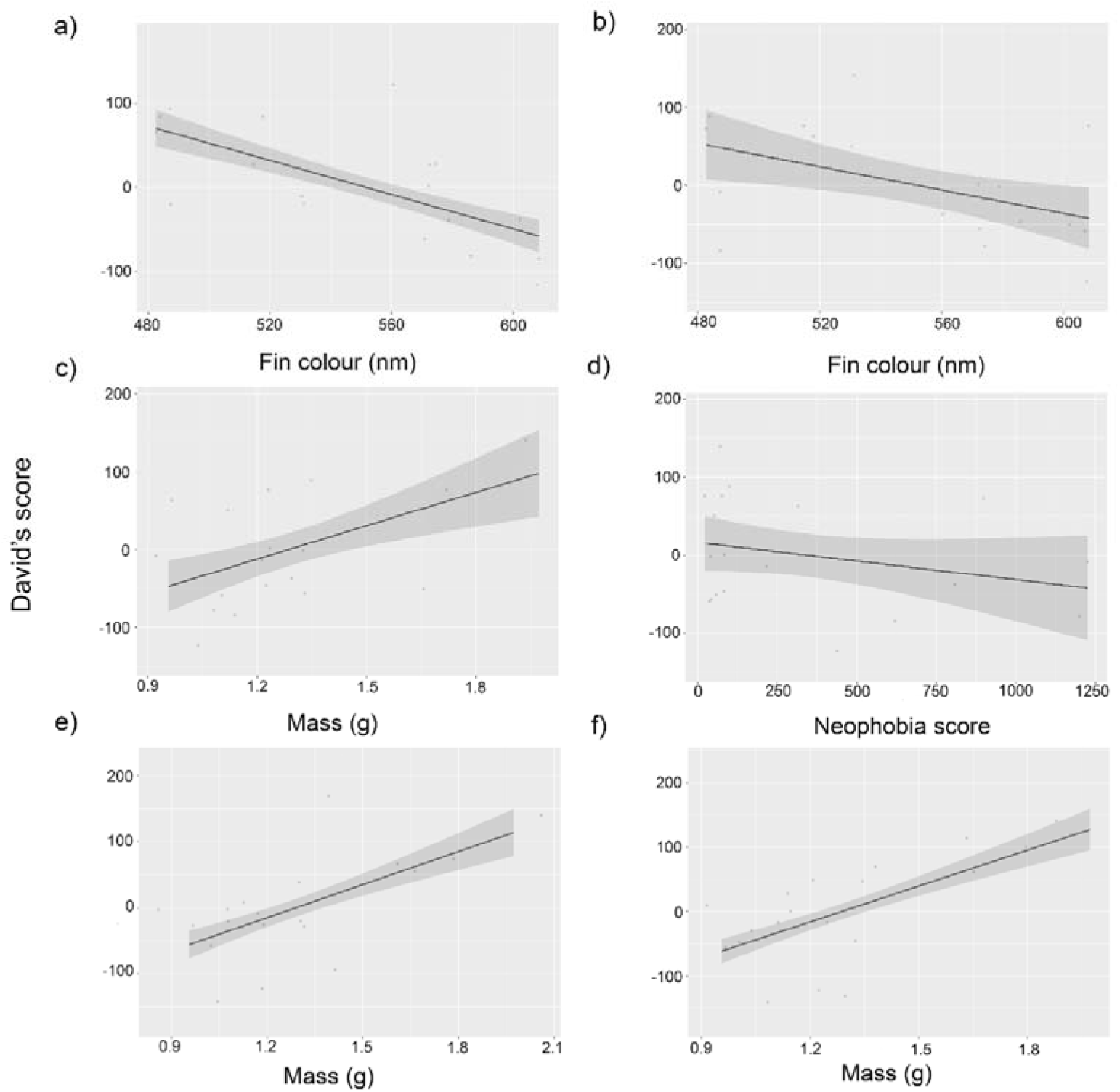
Linear models using David’s score found significant relationships between **a** lateral display duration and fin colour (*P* = 0.03, slope: −0.717), **b** lateral display frequency and fin colour (*P* = 0.007, slope: −0.876), **c** lateral display frequency and mass (*P* = 0.03, slope: 104) **d** lateral display frequency and neophobia (*P* = 0.04, slope: −0.714), e opercular flare duration and body mass (*P* = 0.02, slope: 156), **f** opercular flare frequency and mass (P = 0.01, slope: 172). Confidence intervals are shown in grey.

**Table 1.** The results of the linear models for lateral display duration (DS L_d_) and frequency (DS L_f_) and opercular flare duration (DS O_d_) and frequency (DS O_f_)

| Model | $R^2$ | $F_{4, 14}$ | $P$ |
| --- | --- | --- | --- |
| DS L<sub>d</sub> | 46.5% | 4.91 | 0.012 |
| DS L<sub>f</sub> | 58.0% | 7.22 | 0.002 |
| DS O<sub>d</sub> | 33.0% | 3.21, | 0.046 |
| DS O<sub>f</sub> | 38.5% | 3.81 | 0.027 |
| **DS L<sub>d</sub>** |  |  |  |
| Variable | Slope | Std. error | P value |
| Mass | -15.26 | 45.83 | 0.74 |
| Fin colour | -0.72 | 0.30 | **0.03** |
| Body colour | -0.51 | 0.37 | 0.19 |
| Neophobia | 0.03 | 0.03 | 0.35 |
| **DS L<sub>f</sub>** |  |  |  |
| Variable | Slope | Std. error | P value |
| Mass | 103.60 | 42.71 | **0.03** |
| Fin colour | -0.88 | 0.28 | **0.01** |
| Body colour | -0.34 | 0.35 | 0.34 |
| Neophobia | -0.07 | 0.032 | **0.04** |
| **DS O<sub>d</sub>** |  |  |  |
| Variable | Slope | Std. error | P value |
| Mass | 156.39 | 59.77 | **0.02** |
| Fin colour | -0.66 | 0.39 | 0.12 |
| Body colour | 0.26 | 0.48 | 0.60 |
| Neophobia | -0.04 | 0.05 | 0.38 |
| **DS O<sub>f</sub>** |  |  |  |
| Variable | Slope | Std. error | P value |
| Mass | 171.78 | 58.37 | **0.01** |
| Fin colour | -0.57 | 0.38 | 0.16 |
| Body colour | 0.19 | 0.47 | 0.69 |
| Neophobia | -0.04 | 0.04 | 0.34 |

## DISCUSSION

### Mass

Larger and heavier males tend to be stronger and more aggressive in many species, leading to dominance (Arnott and Elwood 2009; Portugal et al. 2017b). The present study showed that body mass is a predictor of aggression in male *B. splendens*. Mass may also be indicative of fish with a larger labyrinth organ, which could impact opercular flare duration. Opercular flaring concurrently inhibits aquatic respiration for the period that the opercular are raised (Abrahams et al. 2005). As displays between male *B. splendens* are highly intensive and energetically demanding, the increased oxygen requirement (Castro et al. 2006) is met solely by an increase in air-breathing at the surface (Alton et al. 2013). Body mass positively correlates to oxygen uptake per breath (Alton et al. 2013), thus heavier males can afford to perform longer and more frequent opercular flares (Forsatkar et al. 2016).

### Colour

This study indicated that average fin colour is related to aggression, with blue-finned males being most aggressive. Simpson (1968) also demonstrated that blue males displayed more readily and attacked more frequently than different coloured males when shown a mirror or puppet. Since fin flaring is a common behaviour during displays, it is likely that fin colour plays a significant role. Lateral displays are highly visual and males typically display near their opponent (Simpson 1968). Subsequently, aggressive males with a greater fighting ability can risk remaining in close proximity to their opponent.

Like some domestic variations, wild type males have blue fins (Simpson 1968). For wild *B. splendens*, their habitat is resource depleted, hypoxic, and murky (Graham 1997). In these conditions, vision may be limited to a short distance, reducing the effectiveness of visual cues. However, since blue light travels farther underwater (Day 2013), blue signals will be able to reach more opponents and make the colour appear more vibrant. The authors chose not to measure fin length, along with fin clour, as it has previously been shown not to alter aggression in *B. splendens* (Allen and Nicoletto 1997).

### Boldness

Boldness, unlike neophobia, was not found to be repeatable. Neophobic individuals performed more lateral displays than less neophobic individuals. Therefore, neophobic males may adopt a less costly aggressive behaviour to conserve energy and outlast aggressive opponents.

To reduce winner-loser effects altering the outcome and response of males several precautions were put in place. Males were kept in isolation prior to and between encounters to reduce potential effects on the response in following encounters. No individual was tested more than once a day and encounters were randomised to reduce consecutive encounters between the same pairs and habituation. Males were not tested on weekends to provide at least a 60h rest period in social isolation. Additionally, there was a 3-day rest period between each experiment type (aggression, boldness and neophobia) to reduce the studies impacting each other.

In conclusion, analysis of *B. splendens* behavioural and morphological traits revealed several predictors of aggression. Blue-finned males were the most aggressive individuals, performing longer lateral displays, more frequently, while red-finned males were the least aggressive. Additionally, heavier males initiated more opercular and lateral displays, in addition to longer opercular flares. Finally, neophobic individuals were more aggressive, performing more lateral displays. As male *B. splendens* vary in aggression (Simpson 1968), males may adopt different fighting strategies, with some males performing more opercular flares and others adopting a less costly method of fin flaring. Males may have to choose between ending a conflict quickly by escalating to more costly behaviours, such as opercular flaring, or trying to outlast their opponent by conserving energy using less costly behaviours, like fin flaring.

### Ethical approval

All applicable international, national, and/or institutional guidelines for the care and use of animals were followed. The work was approved by Royal Holloway University of London ethics committee.

## ACKNOWLEDGEMENTS

We are grateful to N. Morley and Z. Hale for feeding the fish, M. Tate for his help with observations and The Center for Behavioral Neuroscience for Stopwatch+. This research did not receive any funding from agencies in the public, commercial, or not-for-profit sectors.

